# In silico drug sensitivity predicts subgroup-specific therapeutics in medulloblastoma patients

**DOI:** 10.1101/2025.05.23.655845

**Authors:** Anna M. Jermakowicz, Luz Ruiz, Jonathan Chu, Nitish Jange, Robert K. Suter, Nina S. Kadan-Lottick, Derek Hanson, Nagi G. Ayad

## Abstract

**Background:** Medulloblastoma is the most common malignant pediatric brain tumor. Survival rates vary widely between subgroups, with an average overall survival of 70%. Recurrent medulloblastoma is highly aggressive, treatment-resistant, and usually fatal. In addition, current treatments are highly toxic to the developing brain and surviving patients suffer from lifelong side effects. Therefore, novel therapeutic options are urgently needed.

**Methods:** To inform risk-based, personalized therapy, we developed a novel platform called DrugSeq, which allows predictions of drug sensitivities in patients across medulloblastoma subgroups. We used a perturbagen-response dataset to calculate transcriptional response signatures for each drug and compared this to patient medulloblastoma tumor gene expression. We then stratified patients by molecular subgroup and used an ANOVA analysis to identify drugs that selectively targeted each subgroup.

**Results:** We found distinct differences in transcriptional profiles and predicted drug sensitivity for each medulloblastoma subgroup. We identified several kinase inhibitors, epigenetic inhibitors, and several drugs that have been investigated in drug repositioning studies for cancer.

**Conclusions:** We posit that DrugSeq may identify novel therapies and facilitate patient stratification in clinical trials, leading to more successful targeted medulloblastoma therapies that improve tumor response while minimizing late toxicities. This computational tool can also be used for other cancers to stratify patients based on any clinical or molecular feature.

**Key points:**

⍰ DrugSeq calculates drug sensitivity for medulloblastoma tumors stratified by subgroup.
⍰ DrugSeq platform may inform patient stratification strategies in clinical trials.

**Importance of the Study:** Medulloblastoma is the most common malignant pediatric brain tumor. Current standard-of-care typically includes surgical resection, multi-agent chemotherapy, and radiation. However, survival rates vary widely between subgroups, ranging from 45 to 90%, depending on age and molecular features. In addition, surviving children frequently suffer from debilitating late side effects of therapy including neurocognitive impairment, epilepsy, stroke, subsequent cancer, endocrinopathies, and early mortality. Therefore, novel therapeutic options are urgently needed. However, a one-size-fits-all approach for therapy is unlikely to be effective given the well-characterized intertumor heterogeneity of medulloblastoma.

## Introduction

Medulloblastoma is the most common malignant pediatric brain tumor, accounting for approximately 9% of brain cancer diagnoses in children 0-14 years of age, with an overall 5-year survival rate of 75.5% that varies widely within patient subgroups ^1^. Typically, current treatment consists of maximal safe surgical resection followed by cranial-spinal radiation and multiagent systemic chemotherapy. However in younger children (typically 3-5 years of age and under) radiation is omitted or delayed as long as possible given the particularly severe neurocognitive impairment observed after radiation to the developing brain ^2^.

Medulloblastoma has recently been classified into molecular subgroups based on genomic features and these are the strongest predictors of patient outcomes. These consist of Wingless (WNT), Sonic Hedgehog (SHH), Group 3 (G3), and Group 4 (G4) ^2-5^. Each subgroup has distinct transcriptional profiles, clinical features, prognoses, and treatment outcomes. ^3,6^. Further, these four subgroups have been characterized at the epigenomic and genomic level, allowing for further stratifications within each subgroup ^3,7-9^. At relapse, most tumors retain the same classification as their original subgroup and are often resistant to the previously administered first-line therapy ^5^.

WNT subgroup tumors, account for 10% of all medulloblastomas and are characterized by activating mutations in the WNT/β-catenin signaling pathway ^10^. WNT-activated tumors have the most favorable prognosis, with approximately a 90% survival rate, and relapse is uncommon ^7,11^. The WNT subgroup has been stratified into two subtypes, α and β. The WNT α subtype is mainly diagnosed in children and contains monosomy 6, whereas WNT β is mainly comprised of older patients ^3^.

About 30% of medulloblastoma cases are in the SHH subgroup, which is characterized by dysregulation of SHH signaling. SHH tumors are most common in children under 3 years of age. Younger children with this genetic finding have a better prognosis, with a survival rate of 75% in infants and 50% in children. A disproportionately high proportion of established medulloblastoma mouse models belong to the SHH subgroup as well ^7,12^. There is heterogeneity in SHH subgroups evident in the different age groups including infants, adolescents, and adults ^3,13-15^. The SHH subgroup has been further divided into the subtypes α, β, γ, and δ, each with varying features in histology, gene amplifications, gene deletions, mutations, metastatic dissemination rate, and prognoses ^3^. Among the subtypes, SHH α tumors affect 3-16 year olds and have the worst prognosis. These tumors are enriched with MYCN, GLI2, and YAP1 amplifications. In addition, there is an enrichment of TP53 mutations in SHH α tumors.

While the WNT and SHH subgroups have distinct transcriptional and molecular profiles, G3 and G4 tumors are more difficult to distinguish due to their similar cytogenetic features ^3,7^. G3 medulloblastoma tumors account for approximately 25% of cases. This is the deadliest subgroup of medulloblastoma and the disease is frequently metastatic at the time of diagnosis ^7,11^. These tumors have a poor prognosis and a common feature is MYC amplification. ^9^ Treatment is stratified depending on standard and high-risk G3 groups, which are classified based on metastasis rate, level of resection, and MYC amplification status. ^9^ G3 tumors have been stratified further into three subtypes including α, β, and γ, which helps differentiate G3 from G4 ^3^. There is a more favorable prognosis in G3 α and G3 β tumors compared to G3 γ tumors, which have the worst prognosis ^3^.

G4 is the most common subgroup with approximately an 80% survival rate, and it accounts for 35% of medulloblastoma tumors. These tumors are stratified into G4 α, β, and γ. ^3^ Loss of the X chromosome is common in these tumors, consistent with the high male-to-female ratio of G4 patients ^16^. These tumors are transcriptionally similar to G3 medulloblastoma, however, the current standard of care therapies show distinct differences in overall survival between these groups ^17^.

Upon recurrence, medulloblastoma is often treatment-resistant and fatal, with a five-year survival rate of approximately 25% ^18^. Further, medulloblastoma survivors are frequently burdened with lifelong adverse effects including decreased cognitive abilities, speech difficulties, stunted spinal growth, epilepsy, endocrinopathies, stroke, and even early mortality ^2,19-21^, highlighting the need for novel effective medulloblastoma therapies that are more highly targeted and less toxic.

Despite the extensive molecular characterization of medulloblastoma tumors into distinct subgroups, trials based on molecular biomarkers and subgroups are still limited and are critically needed ^4,22^. There is a diverse response to treatments for each subgroup and subtype, indicating the need to stratify future therapy ^3^. Recent efforts have been made to repurpose existing FDA-approved drugs for cancer treatment since this bypasses the time and cost necessary for drug development and toxicity profiling, allowing more rapid approval for clinical trials ^23^. *In silico* methods have been previously used to identify drugs with desirable predicted target specificity ^24,25^. However, these approaches have not identified potentially patient-specific approaches based on medulloblastoma subgroups. We present a novel means of coupling the transcriptional disease signature of medulloblastoma tumors to compounds that can potentially reverse this signature. To make this pipeline available to the medical and research community, we have created a computational pipeline entitled DrugSeq that will allow patient-specific predictions of the anti-tumor efficacy for both investigational and approved drugs. We posit that DrugSeq will be a useful tool for preclinical and clinical investigations of novel drugs selective for medulloblastoma subgroups.

## Materials and Methods

### Patient gene expression acquisition and processing

Pre-processed microarray or mRNA patient data from Cavalli 2017 and Archer 2018 were obtained ^3,26^. The Cavalli set consists of a total of 763 medulloblastoma samples, consisting of 144 G3 tumors, 326 G4 tumors, 233 SHH tumors, and 71 WNT tumors, and the Archer set consists of 39 medulloblastoma tumors, with 13 G3, 12 G4, and 14 SHH tumors. A pseudocount of 0.01 was added to each transcript expression value and transcripts were averaged by external gene name. Control microarray expression was obtained from Allen Brain Atlas (ABA), using samples from the vermis of the cerebellar cortex ^27^. Cavalli and ABA microarrays were normalized using limma and the log_2_ fold change value for each patient relative to the median control expression was calculated. These patient specific expression profiles, known as the patient disease signatures, were used for downstream analysis.

### Transcriptional consensus scores (TCS)

The Library of Integrated Network-Based Cellular Signatures (LINCS) center at the National Institutes of Health developed a perturbagen-response gene expression profiling platform called L1000. The L1000 assay is a high-throughput bead-based fluorescence assay that measures the steady-state levels of 978 gene transcripts ^28^. The L1000 dataset is a compilation of the transcriptional responses induced by over 1700 drugs across more than 40 cell lines. One previously developed pipeline, SynergySeq, generates a drug-specific transcriptional consensus signature (TCS) containing the L1000 genes that are consistently up or downregulated following treatment with a drug across a multitude of cell lines ^29^. These TCS scores were then used to predict synergy in drug combinations based on the disease transcriptional profile.

### Patient-specific drug response predictions using DrugSeq

To predict the therapeutic response of compounds across medulloblastoma subtypes, we developed a custom R Shiny application that computes transcriptional discordance scores using compound-specific TCSs from L1000 and patient-specific disease signatures. Log_2_ fold change normalized patient gene expression was processed using a novel pipeline called DrugSeq to predict drug sensitivity for each patient (Figure 1). This pipeline uses the previously described compound-specific TCSs and calculates the Spearman correlation between the TCS and the normalized patient gene expression signature, giving us the patient-specific drug discordance. Thus, a more negative score indicates a stronger reversal of the disease signature by the given compound, and therefore the patient is predicted to be more responsive to the drug. The data analysis script is available at https://github.com/AyadLab/DrugSeq and as a user-friendly web portal at https://combin.shinyapps.io/DrugSeq/.

**Figure 1.**
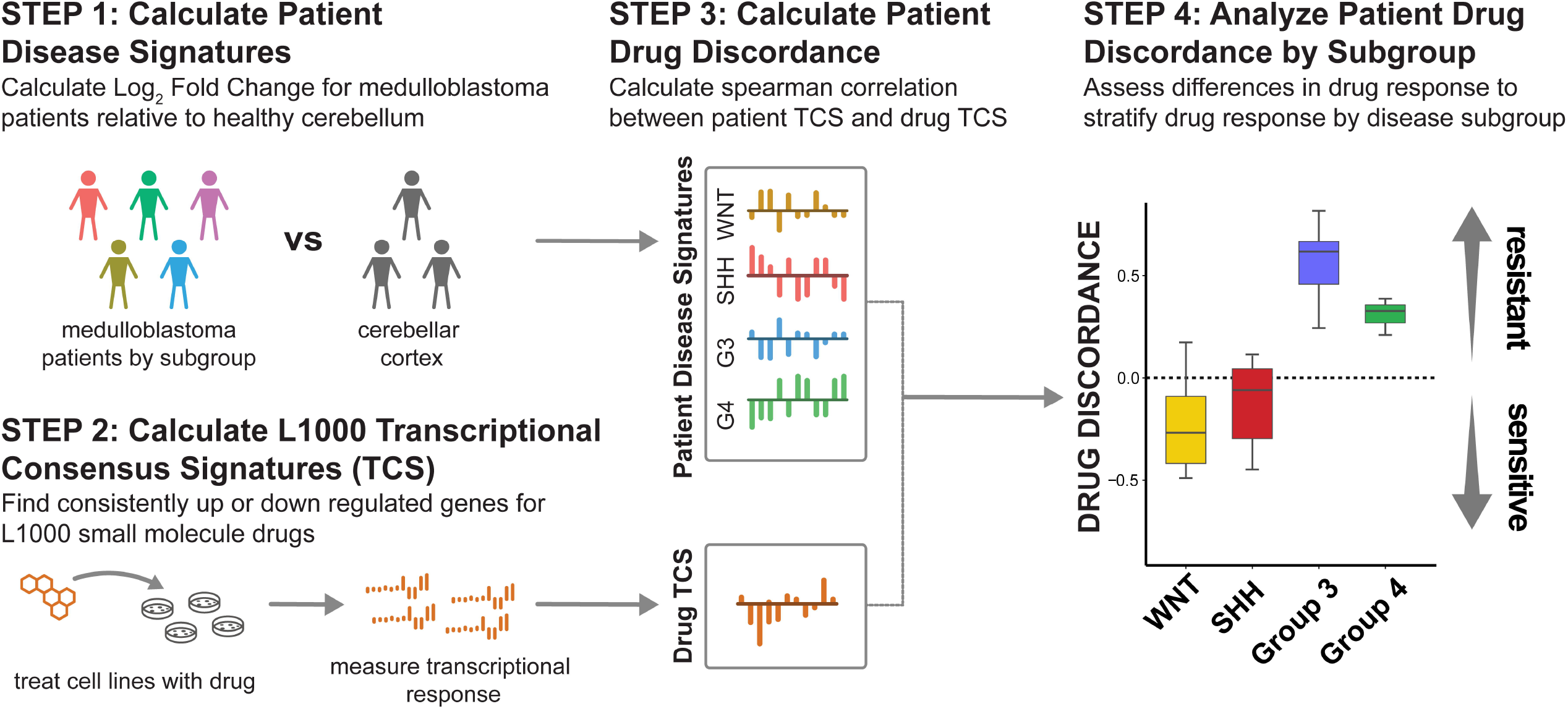
DrugSeq application stratifies individual patient drug response. Patient gene expression is normalized and log_2_ fold change in gene expression is calculated (Step 1). L1000 transcriptional consensus signatures (TCS) are calculated (Step 2) and discordance between the disease signature and the drug signature is calculated for each patient (Step 3). Patients are stratified by subgroup and differences in drug discordance by subgroup are calculated using an ANOVA with Tukey HSD correction (Step 4).

### Subgroup-specific DrugSeq sensitivity predictions

To identify therapies that are predicted to target tumor cells in a subgroup-specific manner, we performed an ANOVA analysis with Tukey HSD test on the patient-specific drug discordance scores for each drug. Patients were stratified by subgroups, including G3, G4, SHH, and WNT medulloblastoma. A one-way ANOVA test was used to find drugs that vary in patient response by subgroup, then a Tukey HSD test was used to determine which subgroups varied. A separation score was calculated to quantify the degree of separation in predicted response between groups relative to within-group variability. This was defined as the difference between the highest and lowest subtype mean responses, divided by the average within-subtype standard deviation (pooled SD). Results were filtered by minimum size of TCS (≥ 20 genes), predicted blood-brain barrier permeability (logBB > -1), predicted sensitivity (minimum response across all groups < 0). Among these, the most sensitive subtype for the drug was identified and drugs were ranked by descending separation score. The top twenty compounds were selected by separation score and plotted on a heatmap (Figure 2A-B).

**Figure 2.**
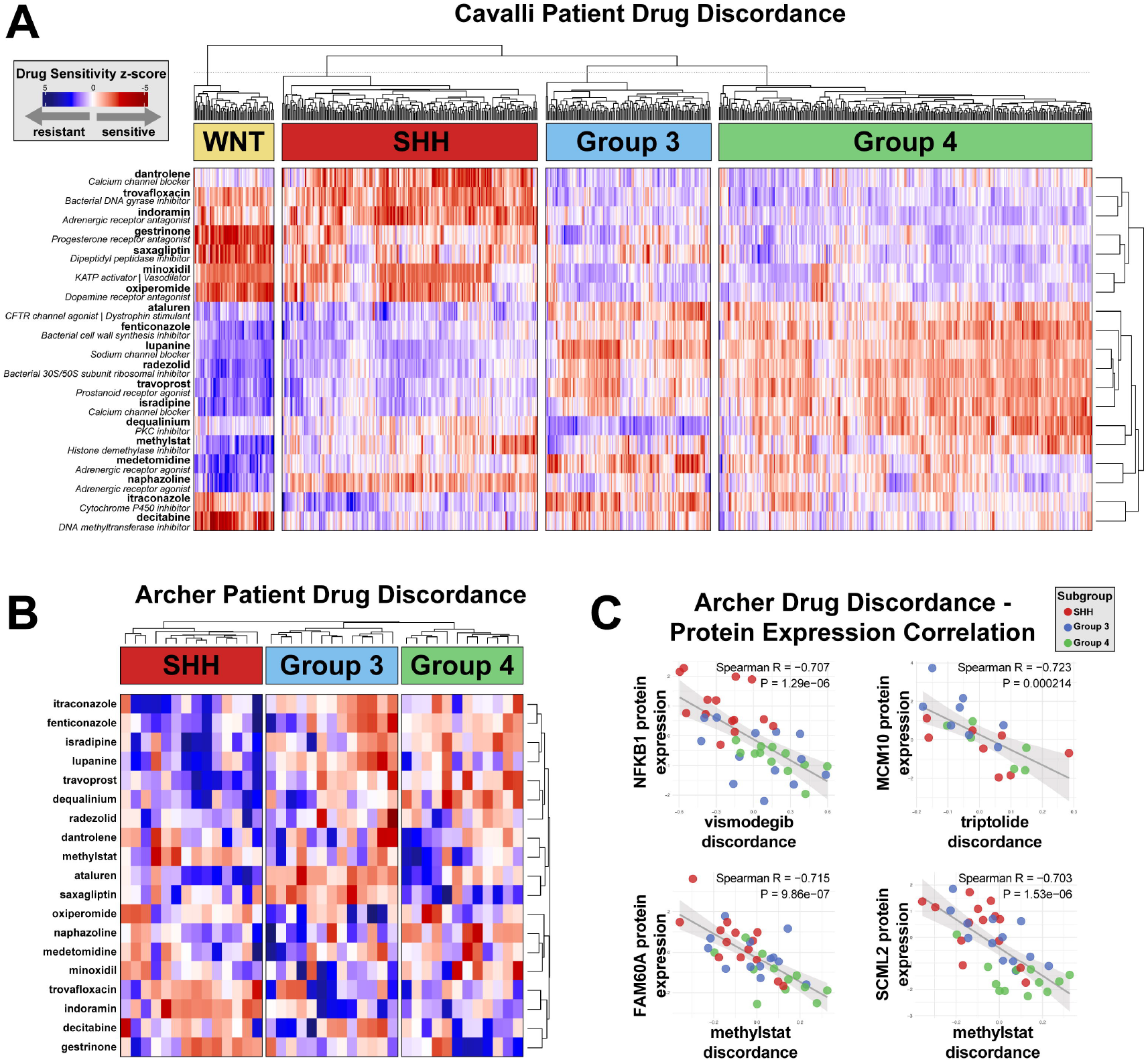
Patient-specific drug discordance reveals distinct drug sensitivities in medulloblastoma subgroup. **A. Heatmap of top hits by subgroup.** DrugSeq drug discordance scores were calculated for each patient. Scores were filtered for drugs that were a hit in any subgroup (minimum mean discordance > 0), and the top anti-cancer drugs were selected using a one-way ANOVA test. **B. Heatmap of top hits for Archer dataset**. Drug discordance scores for top inhibitors were plotted with patients separated by medulloblastoma subgroup or recurrence. **C. Identification of protein expression correlated with drug discordance for top compounds**. Patient-matched protein expression from the Archer dataset was correlated with predicted drug sensitivity. Significant correlations between drug target protein expression and predicted drug sensitivity for selected drugs are shown.

### Patient protein expression

Protein expression from the Archer 2018 cohort was obtained and normalized to the average expression across all samples for each protein. Protein targets for top inhibitors were identified using PubChem and correlations between target protein expression and drug discordance was calculated (Figure 2C).

### Mechanism of action enrichment analysis

To determine enriched mechanisms of actions for each subgroup, a list of subgroup- or subtype-specific drugs was generated by filtering for compounds that had the highest mean discordance for each group. A comprehensive list of all MOAs for all compounds was curated and the frequency of each MOA was calculated across the entire set of drugs to establish a background frequency. Next, for each group drugs were filtered for subgroup-specificity where only those with a predicted mean discordance <-0.5 to ensure meaningful differential sensitivity. These group-specific drugs were then joined with MOA annotations to establish a group-specific frequency for each MOA. Next, the group-specific frequency was divided by the background frequency for each MOA to calculate the percentage of enrichment for the MOA. Finally, the percentage enrichment was multiplied by the count per group to weight MOAs occurring multiple times in the group. For each subgroup or subtype, the top 10 enriched MOAs were visualized on bar plots (Figure 3, Supplementary Figure 1).

**Figure 3.**
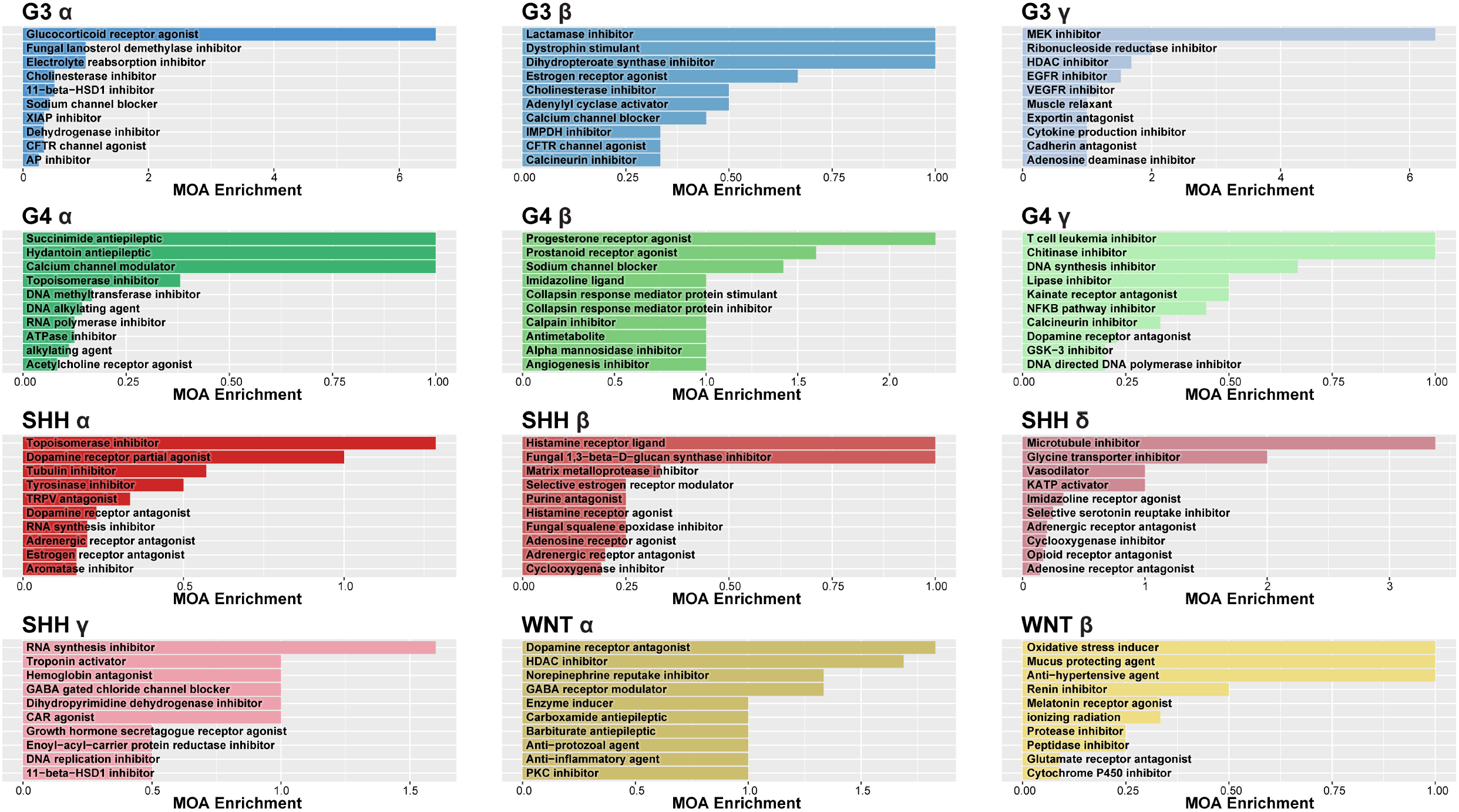
Medulloblastoma subgroups have distinct drug targets. DrugSeq drug discordance scores were calculated for each patient in the Cavalli dataset. Top compounds for each subtype were determined by filtering for the most discordant subgroup for each compound. Compounds were compiled based on MOA to determine enrichment of drug targets within the top hits.

### *In vitro* drug response

S47 and GFAP-Cre;Ptc1fl/fl are SHH-activated medulloblastoma cell lines derived from mice, and MB03 and D425 cells are derived from human Group 3 medulloblastoma tumors. Of note, S47, GFAP-Cre;Ptch1fl/fl, and D425 cells are grown in suspension, while MB03 cells are adherent. We treated cells with 1:2 serial dilutions of methylstat and measured the ATP content via CellTiter-Glo after a 3-day incubation. Response was normalized to DMSO as a negative control and 10uM velcade as a positive control (Figure 4A).

**Figure 4.**
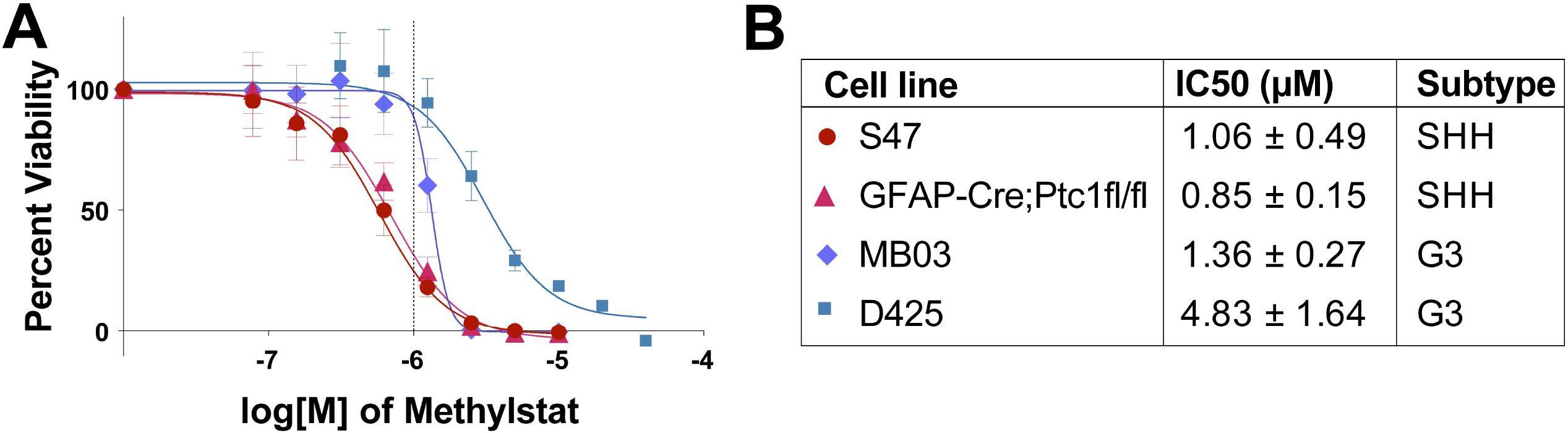
Top SHH inhibitor related drug induces cell death in vitro. **A. Methylstat reduces viability in SHH medulloblastoma cells.** SHH-MB cell lines (S47 and Tg(GFAP-Cre)) and G3-MB cell lines (MB03 and D425) were treated with 1:2 dilutions of methylstat. **A. Dose response curve for methylstat in MB cells**. Plot displayed is representative of 3 biological replicates (8 technical replicates). **B. SHH-MB cells are more sensitive to methylstat**. IC50 values were calculated for each cell line.

### Drug and radiation synergy in medulloblastoma

To evaluate possible combination therapies with medulloblastoma drugs, we used the previously developed pipeline SynergySeq^29^. Briefly, this pipeline takes the normalized patient disease signature and calculates the disease reversal to the first drug to be used in synergy, then ranks the remaining drugs by disease signature reversal.

#### Combining monotherapy and combination therapy predictions

To evaluate the ability of our novel platform to predict drug response compared to the established SynergySeq platform, we analyzed drug sensitivity across recurrent medulloblastoma patients using our DrugSeq discordance scores and compared those results to SynergySeq predictions for radiation synergy in newly diagnosed patients. For this analysis, we used a dataset from the Pediatric Brain Tumor Atlas (PBTA) from PedcBioPortal, including a total of 81 medulloblastoma samples (7 Group 3 tumors, 44 Group 4 tumors, 24 SHH tumors, 3 WNT tumors, and 8 recurrent tumors). We plotted the average drug discordance score for each drug and calculated the Spearman correlation between the DrugSeq predicted response in recurrent patients (previously treated with radiation therapy) to newly diagnosed (untreated) patients (Figure 5A).

**Figure 5.**
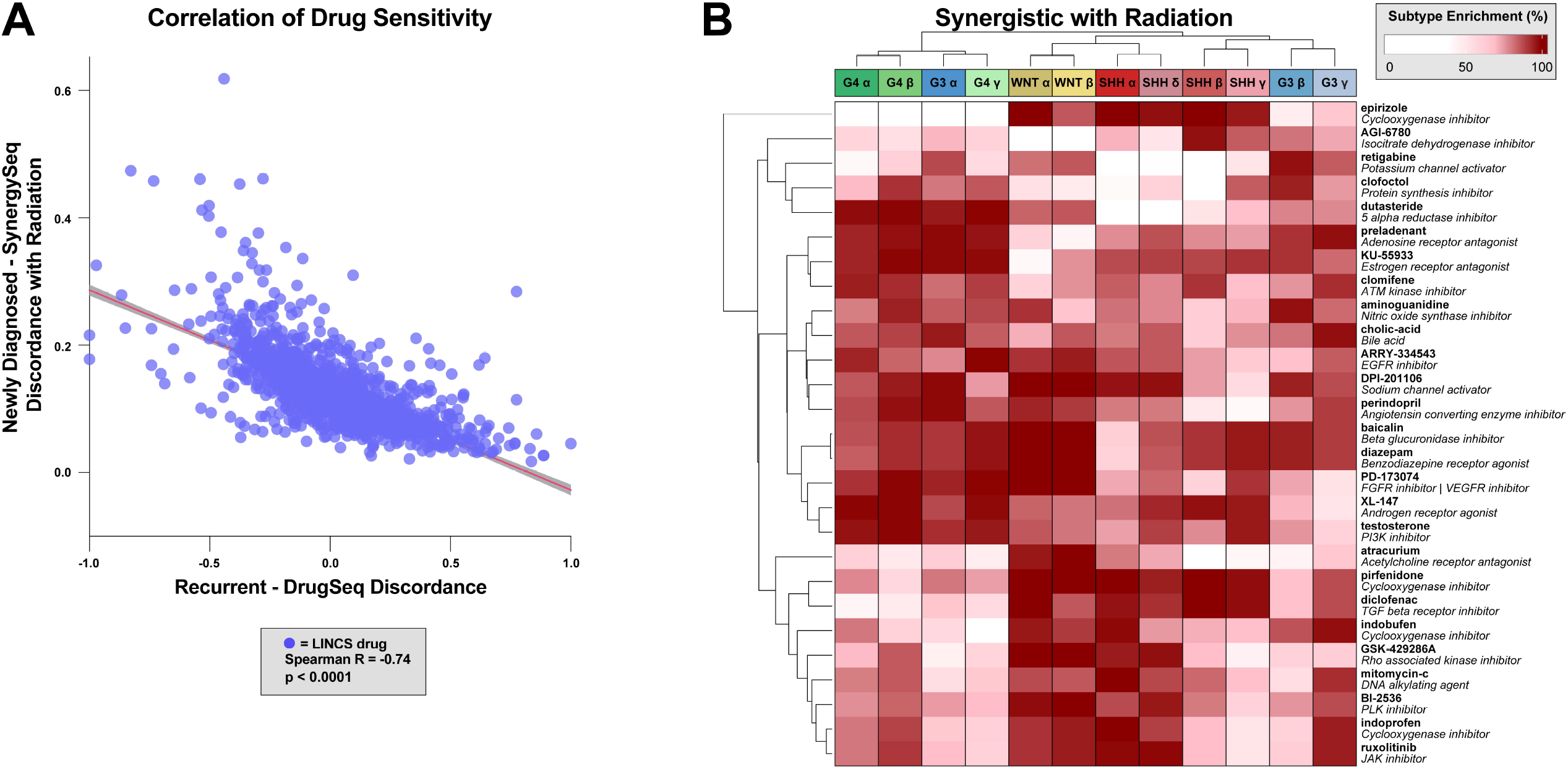
Synergistic combination therapy with radiotherapy in newly diagnosed medulloblastoma tumors correlates with monotherapy discordance in recurrent medulloblastoma. **A. DrugSeq and SynergySeq predictions are significantly correlated.** The average drug discordance score for each drug either in monotherapy for recurrent tumors (previously irradiated) or combination therapy for newly diagnosed tumors (radiation naïve) was plotted and the Spearman correlation between the two cohorts was calculated. **B. SynergySeq reveals drugs that are synergistic with radiation therapy between subgroups**. Individual patient SynergySeq LOS scores were calculated for each drug in combination with radiation. Enrichment within each subgroup was calculated for each inhibitor and the top hits were plotted on a heatmap.

#### Drug synergy predictions with radiation therapy

We calculated the LINCS Orthogonality Scores (LOS) for each patient in combination therapy with radiation therapy. LOS scores greater than one were considered to be synergistic, and the proportion of patients across each subtype that were predicted to have a synergistic response to each inhibitor was calculated. To determine combination therapies that would be differentially effective across subgroups and subtypes, we used a one-way ANOVA test to determine significant differences in LOS scores across subtypes. We filtered for ANOVA < 0.05, subgroup enrichment > 0.75 in at least one group, and anti-cancer drug activity. We then selected the top four combination drugs for each reference drug and plotted the subtype enrichment of patients with predicted synergy for each subgroup (Figure 5B).

## Results

### DrugSeq platform identifies drug sensitivity by subgroup of medulloblastoma

The DrugSeq platform was used to identify drugs that have differential sensitivity across medulloblastoma subgroups (Figures 1-3). Consistent with the gene expression profiles, G3 and G4 tumors showed the most overlap in predicted drug sensitivity across the subgroups, while SHH and WNT tumors shared similar drug sensitivity profiles (Figure 2A-B, Figure 3). As a sanity check, several known medulloblastoma therapies were identified and subgroup-specific sensitivity was assessed. As expected, SHH-MB tumors were found to be most sensitive to the hedgehog pathway inhibitors vismodegib and taladegib (Supplementary Figure 2A-B)^30,31^. Triptolide, which was recently described in Group 3-MB was found to be most sensitive for Group 3 and Group 4 tumors (Supplementary Figure 2C)^32^. Finally, radiation therapy was found to be most effective in WNT-MB, which is consistent with the literature (Supplementary Figure 2D)^33,34^. Next, drugs displaying sensitivity across all subgroups were analyzed based on mechanism of action (MOA) to determine overlapping and specific MOAs (Figure 3, Supplementary Figure 1). This showed a high degree of overlap for drug targets across all subgroups, as well as specific vulnerabilities with each subgroup.

#### WNT-activated drug sensitivities

Tolvaptan, a vasopressin V2 receptor antagonist, has been shown to reduce neuroblastoma growth in vitro ^35^. Decitabine, a DNA methyltransferase inhibitor, has been evaluated in combination with chemotherapy, radiotherapy, and immunotherapy ^36,37^. Bruceantin, a protein synthesis inhibitor, has been shown to have antitumor activity in multiple cancers^38^. The effects of bruceantin on c-MYC expression may make it a promising therapeutic option for WNT-activated MB. I-BET-151, a bromodomain inhibitor, also suppresses MYC expression and may downregulate WNT signaling. EMD_1214063 (tepotinib) has also been shown to inhibit tumor growth by targeting the WNT pathway^39^. Mechanisms of action identified as specific for WNT activated tumors included proteasome inhibitors and DNA replication inhibitors^40^.

#### SHH-activated drug sensitivities

Thioguanine, a purine analog, is FDA-approved for AML treatment. Further, it has been shown to inhibit GLI2, which is part of the SHH-pathway ^41^. Methylstat, a selective inhibitor of Jumonji C domain-containing histone demethylase, has been shown to inhibit angiogenesis in several cancer types including multiple myeloma ^42,43^. JMJD3, a JMJC component, has been shown to be activated by SHH signaling in medulloblastoma ^43^. Mechanisms of action specific for SHH-driven MB included adenosine receptor antagonists and purine antagonists.

#### Group 3 drug sensitivities

Among the top inhibitors were NVP-AEW541, cladribine, birinapant, and ataluren. NVP-AEW541, an IGF-IR kinase inhibitor, may be a promising treatment for medulloblastoma. IGF-IR has been shown to be activated and overexpressed in medulloblastoma, contributing to tumor growth^44^. Cladribine, a purine analog, has shown promise in hematological malignancies, as well as non-WNT activated MB ^45^. The IAP antagonist birinapant has been shown to synergize with CAR T-cell therapy in glioblastoma, and radiosensitize tumors^46^. Ataluren, a drug developed for the treatment of muscular dystrophy, promotes read-through of premature stop codons, which could provide a therapeutic benefit for tumors with mutations in tumor suppressor genes^47^. Ataluren is undergoing a clinical trial for colorectal cancer for synergy with an immune checkpoint inhibitor [NCT04014530].

#### Group 4 drug sensitivities

JNK-9L is a JNK inhibitor that has been shown to be brain penetrant *in vivo* ^48^. Novobiocin, traditionally an antibiotic, has been shown to target DNA polymerase theta and can target some types of cancer, including glioblastoma ^49^. Further, thiabendazole, an antifungal medication, has been shown to target MCM2 in glioblastoma cells, limiting proliferation and invasion ^50^. Remarkably, MOAs identified for Group 3 and Group 4 medulloblastoma displayed a very high degree of overlap, highlighting the similarities between these subgroups (Supplementary Figure 1).

### Drug sensitivity correlates with protein expression to provide insight into drivers of drug response

To investigate the potential drivers of drug response in medulloblastoma, we explored the correlation of predicted drug sensitivity and protein expression across a panel of patients. Using proteomic profiles and drug sensitivity predictions, we performed correlation analyses to identify proteins whose expression levels were significantly associated with response to methylstat. Two proteins, FAM60A and SCML2, were found to be strongly and significantly correlated with drug sensitivity (Figure 2C). High expression of both proteins was associated with increased sensitivity to methylstat across the patient cohort. These proteins are both part of complexes that regulate histone methylation ^51,52^, suggesting a potential mechanistic link to the drug’s mode of action.

### *In vitro* cell viability reflects subgroup-specific drug sensitivity predictions

As a proof of concept, we tested the compound methylstat in a panel of medulloblastoma cell lines, including two SHH-activated cell lines (S47 and Tg(GFAP-Cre)) and two Group 3 cell lines (MB03 and Ptcfl/fl). We found that SHH-activated cell lines had reduced IC50 and reduced viability of in vitro, compared to Group 3 cell lines (Figure 4A). The IC50 values for S47 and Tg(GFAP-Cre) (1.06, 0.85 µM) were significantly lower than those for MB03 and Ptcfl/fl (1.36, 4.83 µM), indicating increased sensitivity to methylstat (Figure 4B). Cell viability assays showed a dose-dependent decrease in proliferation in all cell lines, with over 80% reduction at 1 µM. By contrast, Group 3 cell lines exhibited modest changes in viability, with only 25% reduction in viability at 1 µM (Figure 4A, dashed line).

### Drug sensitivity for recurrent patients correlates with predicted synergy with radiation in newly diagnosed patients

To assess the ability of the DrugSeq platform to predict drug sensitivity with radiation therapy, we looked at the correlation between predicted drug sensitivity in recurrent patients and drug synergy with radiation in newly diagnosed patients. To do this, we used the disease discordance scores from the SynergySeq predictions with radiation in newly diagnosed patients and the DrugSeq drug discordance score in recurrent patients. Discordance scores were averaged across all patients, and a Spearman correlation was performed. We found a strong correlation between these drug sensitivity predictions, with a Spearman R of -0.74 and a P-value < 0.0001, suggesting that DrugSeq may be a valuable tool for determining synergistic combinations with radiotherapy (Figure 5A).

### Synergistic drug responses with radiation across medulloblastoma subtypes

Drug synergy with radiation therapy was evaluated across the twelve molecular subtypes of MB, as determined by Cavalli et al. ^3^. Hierarchical clustering of drugs reveals distinct patterns of synergy. Several cyclooxygenase inhibitors (epirizole, diclofenac, indobufen, and indoprefen) displayed strong synergy in Group 3 and Group 4 subtypes. This suggests a potential vulnerability in these subtypes to combined inhibition of inflammatory pathways and radiation, which is consistent with literature demonstrating the radiosensitizing and radioprotective effects of COX-2 inhibitors ^53,54^. SHH subtypes exhibited enrichment for DNA alkylating agents (mitomycin C) and kinase inhibitors (BI-2536). Mitomycin C has been evaluated in clinical trials with positive results as adjunct to radiation therapy ^55^. BI-2536, a PLK1 inhibitor, has been evaluated for combination with radiation in SHH-MB with positive preclinical results^14^. Collectively, these results underscore the potential for subtype-specific therapies in combination with conventional therapy to enhance treatment efficacy (Figure 5B).

## Discussion

We report distinct drug sensitivity profiles across medulloblastoma subgroups. We developed a platform called DrugSeq, which is freely available for use at https://combin.shinyapps.io/DrugSeq. DrugSeq users can upload a panel of patient gene expression profiles with associated metadata subgroups to assess predicted drug sensitivity (Figure 6A). DrugSeq compares the patient transcriptional profile to the L1000 perturbagen-response transcriptional consensus scores and predicts drug sensitivity for each individual patient (Figure 6B). Using an ANOVA test with Tukey HSD, DrugSeq then calculates drugs that vary significantly across groups of patients (Figure 6C). Finally, users can generate boxplots of drug sensitivity scores for drugs of interest separated by subgroups (Figure 6D).

**Figure 6.**
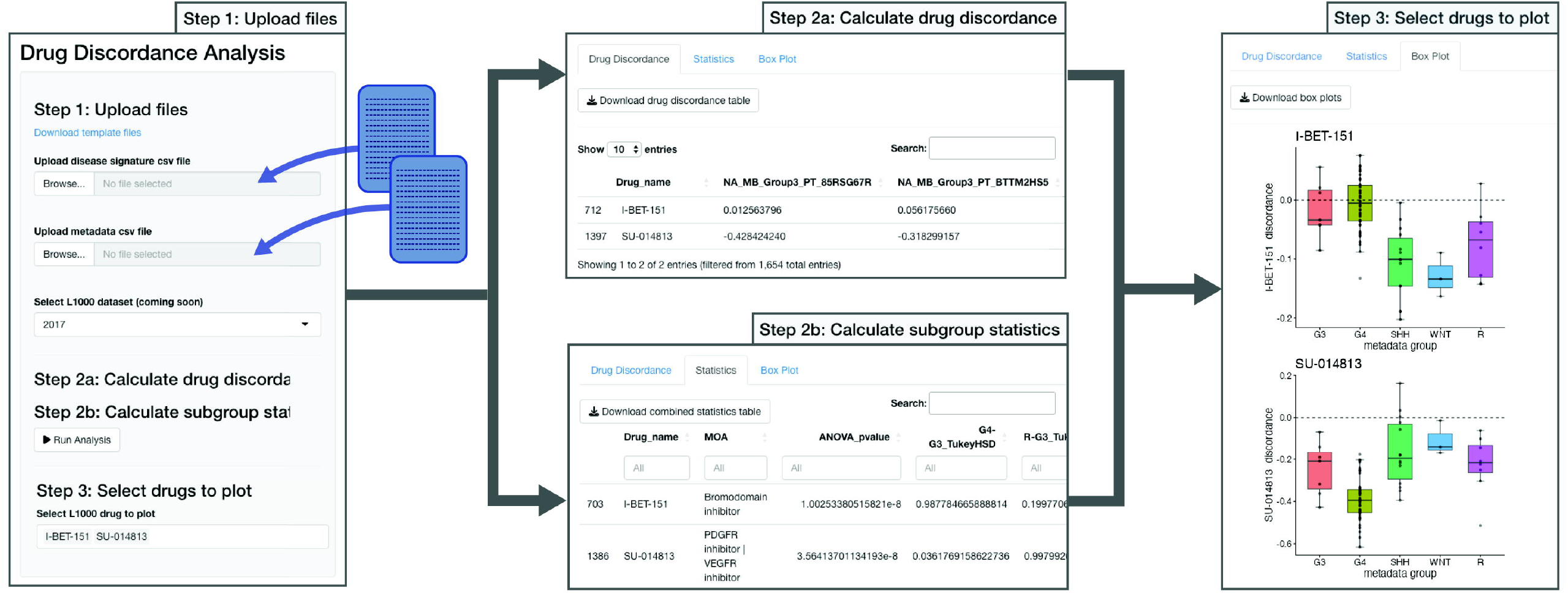
DrugSeq web portal. **A**. User uploads disease signature and metadata files, then selects the L1000 drug dataset to analyze. **B-C**. DrugSeq calculates the drug discordance for each patient (B) and statistics across subgroups (C) for each drug. **D**. User selects drugs to visualize on boxplots.

For our analysis of medulloblastoma patient tumors, we identified compounds that were predicted to target patients in distinct molecular subgroups. We found that G3 and G4 medulloblastoma tumors shared similar profiles of drug sensitivity, consistent with their similar transcriptional profiles. Our data provide valuable information that is critical for stratifying patients into treatment groups for clinical trials. Using this platform, we can predict subgroup-specific and even patient-specific drug vulnerabilities in cancers.

We found that G3 specific tumors are predicted to be targeted by triptolide, which we have previously demonstrated to preferentially inhibit G3 dependent tumor growth in vitro and in vivo ^32^. This provides further validation that predictions from our DrugSeq pipeline are confirmed both in vitro and in vivo in animal models of MB. Indeed, our novel studies here suggest that methylstat, which is predicted to target SHH-MB preferentially induces heightened cell killing of SHH-subtype cells in vitro and ex vivo. Collectively, these studies suggest that there is a significant correlation between what we predict using DrugSeq and what is seen in vitro or in animal models of MB.

Drug repositioning is a powerful tool that can be used to fast-track drug development for brain cancers. This process involves taking an already FDA-approved drug, or a drug in advanced clinical trials, and expanding the treatment criteria to include cancers such as medulloblastoma.

This is advantageous because safety profiles and dosing regimens are already established for both animal studies and eventual clinical trials. Furthermore, brain penetrance and toxicities are often already determined. Prior drug repositioning studies have relied heavily on observational data, such as associations between patient survival and previously prescribed medication for cardiovascular diseases ^56,57^, as well as large *in vitro* drug screens leading to discoveries such as SHH pathway components being targeted by antifungal drugs ^58,59^. By contrast, *in silico* approaches to drug repositioning often rely on structure-based modeling to identify drugs that target a known oncogene, for instance targeting smoothened (SMO), the positive regulator of the hedgehog pathway, for SHH subgroup tumors in medulloblastoma. However, these inhibitors have often failed in the clinic due to downstream mutations in the SMO pathway and acquired resistance ^49^. Other *in silico* models, such as ours, focus on finding new mechanisms of action for known drugs using transcriptional profiles ^60-62^.

Using the DrugSeq pipeline, drugs can then be filtered by known anti-cancer relevance, blood brain barrier permeability, or mechanism of action. For instance, psychiatric drugs have frequently been found to be effective at inhibiting brain tumor cell growth ^63-65^. Elevated dopamine levels, which are achieved with drugs such as butaclamol, have been shown to improve the efficacy of chemotherapeutic agents ^66^. We were also able to predict synergistic combinations, suggesting potential combination therapy with decitabine, a hypomethylating agent, and radiation in WNT, Group 3, and Group 4 tumors. Furthermore, we were able to compare compounds predicted to target recurrent tumors that were previously irradiated and compare these to compound synergy predictions with radiation in newly diagnosed (non-irradiated) tumors. We found a strong correlation between the drug predictions, suggesting that DrugSeq may provide a way to predict drug sensitivity in recurrent medulloblastoma, as well as in combination with radiation therapy.

Large-scale drug screening assays to assess drug sensitivity in vivo are less powerful for medulloblastoma since the majority of cell and animal models of medulloblastoma are representative of the SHH subgroup ^67^. We believe that DrugSeq can help bridge this gap in screening by prioritizing drugs for distinct transcriptional subgroups of medulloblastoma, allowing clinicians to stratify patients in clinical trials. Further preclinical studies of drug candidates will need to be performed *in vitro* and *in vivo* to assess the effect on tumor growth.

In conclusion, we have developed a powerful computational platform that allows drug sensitivity predictions for individual patients, paired with statistical analysis to identify drugs that target subsets of patients. This pipeline can be used for additional cancer types and various subgroups, allowing predictions of drug sensitivities for various features in cancer patients.

## Acknowledgments

We would like to thank all members of the Ayad Lab for the helpful discussion and editing of this manuscript, including. Simon Kaeppeli, Jordan Walter, Matthew D’Antuono, and Emma Rowland.

