## Supplementary Figure for "In silico drug sensitivity predicts subgroup-specific therapeutics in medulloblastoma patients"

**Supplementary Figures**

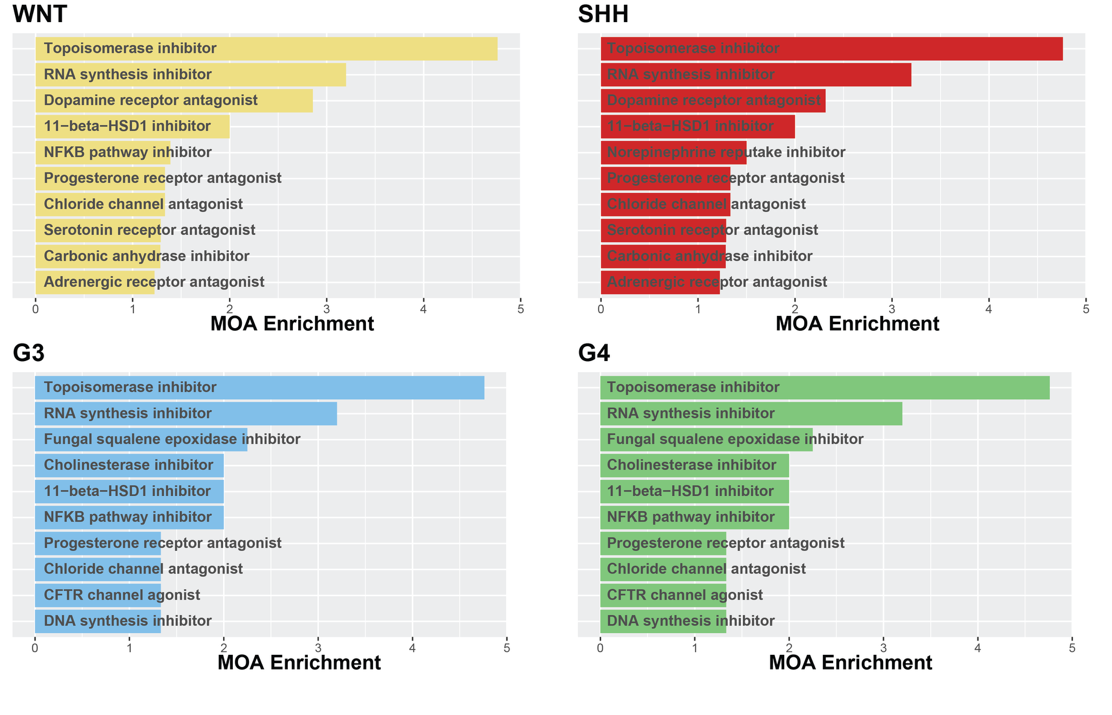

**Supplementary Figure 1. Medulloblastoma subgroups have overlapping drug targets.** DrugSeq drug discordance scores were calculated for each patient in the Cavalli dataset. Top compounds for each subgroup were determined by assessing all compounds with predicted sensitivity. Compounds were compiled based on MOA to determine enrichment of drug targets within the top hits.

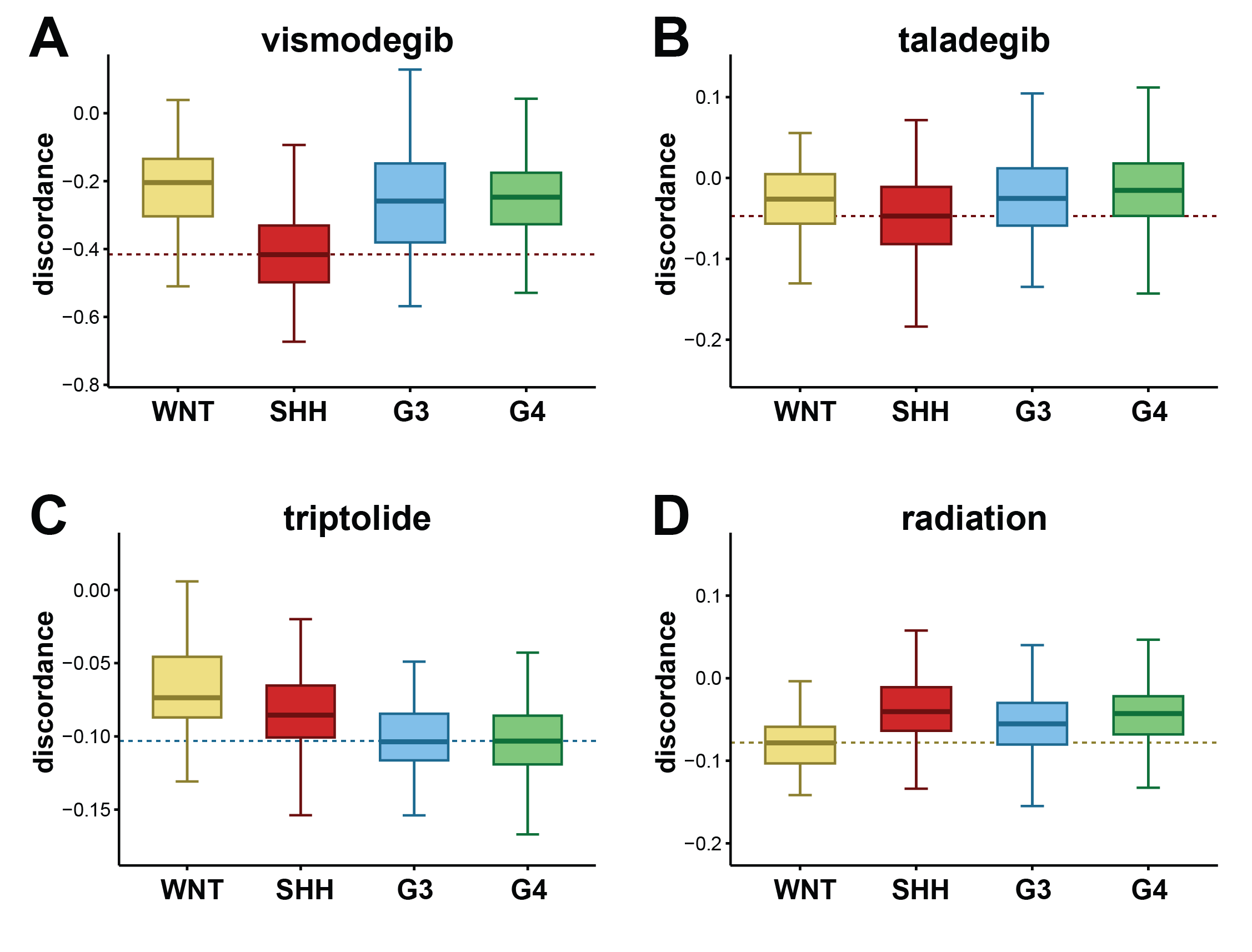

**Supplementary Figure 2.** DrugSeq drug discordance scores were calculated for each patient and known medulloblastoma therapies were visualized on a boxplot. Dashed lines indicate minimum median subgroup sensitivity. **A-B.** Hedgehog pathway inhibitors vismodegib and taladegib. **C.** Group 3 inhibitor triptolide. **D.** WNT-MB inhibitor radiation therapy.
